# Metabolic Control Analysis of Exponential Growth and Product Formation

**DOI:** 10.1101/485680

**Authors:** David A. Fell

## Abstract

Metabolic Control Analysis defines the relationships between the change in activity of an enzyme and the resulting impacts on metabolic fluxes and metabolite concentrations at steady state. In many biotechnological applications of metabolic engineering, however, the goal is to alter the product yield. In this case, although metabolism may be at a pseudo–steady state, the amount of biomass catalysing the metabolism can be growing exponentially. Here, expressions are derived that relate the change in activity of an enzyme and its flux control coefficient to the change in yield from an exponentially growing system. Conversely, the expressions allow estimation of an enzyme’s flux control coefficient over the pathway generating the product from measurements of the changes in enzyme activity and yield.

## 1 Introduction

Metabolic Control Analysis (MCA) (Kacser and Burns; 1973; Heinrich and Rapoport; 1974; Fell; 1997) was developed as a theory to underpin understanding of metabolic control and regulation. For metabolic pathways at steady state, it characterizes the relationships between an alteration of enzyme activity, however caused, and the resulting effects on metabolic fluxes and metabolite concentrations. It has also been extended to certain time– dependent phenomena including transitions to a new steady state after a perturbation (e.g. Acerenza et al.; 1989; Meléndez-Hevia et al.; 1990). In terms of application to metabolic engineering, a limitation of the theory is that it is a logarithmic approximation close to the steady state, and loses accuracy for the large changes in activity induced by over–expression of enzymes since the control coefficients defined in MCA themselves vary as enzyme activities change. However, in many of the cases studied experimentally, the value of the flux control coefficient varies as an approximate hyperbolic function of the enzyme activity; see Kacser and Burns (1973) and Fell (1997) for examples. In such cases, Small and Kacser (1993a,b) derived relationships relating large changes in enzyme activity and the corresponding changes in flux to the values of the conventional flux control coefficients in the original and perturbed states.

Though the finite change relationships of Small and Kacser (1993a) combined with a knowledge of flux control coefficients can be used to design metabolic engineering applications, the desired goal is generally not just to obtain better flux to the target product but to obtain an increase in yield at the end of the process. In many cases, such as batch fermentations of microbial cells or the filling of seed embryos of brassicas with oil, the metabolically active biomass is increasing exponentially throughout the process, and though metabolism remains in a pseudo-steady state, the yield is a time–dependent process and is not directly predicted by the finite change relationships. Here I develop the equations to describe how engineering an altered enzyme activity affects the yield and the relationship with the flux control coefficient.

## 2 Basic functions and definitions

Exponential growth is defined by the differential equation for the rate of growth:

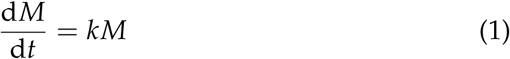

where *k* is the rate constant and *M* is the amount (or concentration) of biomass, or of a product produced in constant proportion to the biomass in the exponential growth phase. For incorporation into Metabolic Control Analysis, we note that d*M*/d*t* is termed a flux, *J*, in that context.

The aim of the derivations below is to extend the explicit equations that describe the impacct of altering an enzyme’s activity on fluxes and concentrations of a biochemical system at steady state to the case where the system is growing exponentially.

A central concern of MCA is to provide an explicit analysis of the impact of changing an enzyme’s activity, such as with a specific inhibitor or by altering its expression level, on the steady state values of variables of the system, such as the metabolic fluxes *J*. The measure used for this is the flux control coefficient, defined as:

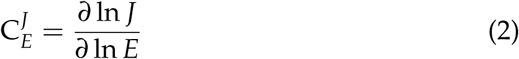

where *E* is the amount or activity of the chosen enzyme *E*. The logarithmic formulation implies that 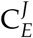 is normalised so that it indicates the fractional change in *J* for the fractional change in *E* (or popularly, the percentage change in *J* for a 1% change in *E*).

Since *J* = *kM*, for any given value of *M* we can write:

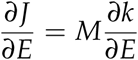

Multiplying both sides by *E*/*J* and substituting for *J* on the RHS by *kM* gives:

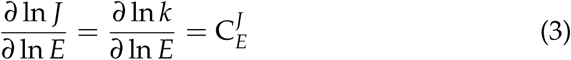

In other words, as might have been expected, the control coefficient describing the effect of *E* on the rate constant *k*, 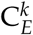, is identical to the flux control coefficient.

Finally, note that the definition of the flux control coefficient can be extended to characterise the effect of a change in enzyme activity on any variable *V* of the metabolic system at steady state as 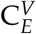.

## 3 Reponse of a growing system as a function of time

The solution of Eqn. 1 is:

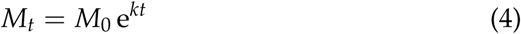

where *M* = *M*_0_ at *t* = 0. Here we want to determine the response of *M*_1_ to a change in activity of a specific enzyme *E* from a constant starting state of the system *M*_0_. Differentiating with respect to *E* gives:

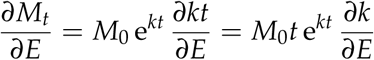

Multiplying by *E*/*k* and substituting *M_t_* for *M*_0_ e*^kt^* gives:

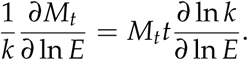

Rearranging gives the final expression for the response 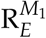:

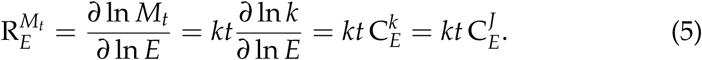

This shows that the response of the perturbed system increasingly diverges from the reference as a function of time, and amplifies the effect of 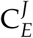 for values of *kt* > 1. Equivalently, this response coefficient is a measure of the divergence between the value of *M_t_* in the unperturbed system (*M*_*t*,*E*_) and the perturbed one (*M*_*t*,*E*+*δE*_), both of which are a function of time.

The actual value of *M*_*t*,*E*+*δE*_ after a perturbation of enzyme activity *δ E*/*E* is obtained by first using the response function to compute the resulting *δ* ln *M_t_*:

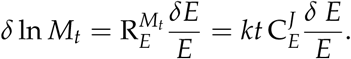

Adding this to the expression for ln *M_t_* from the logarithmic form of Eqn. 4 gives:

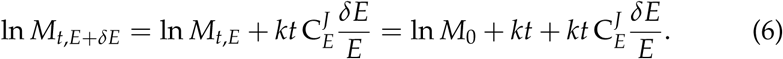

Collecting terms and reverting to exponential format gives:

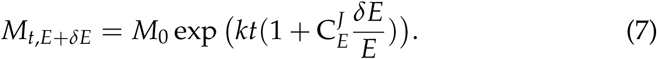

Also from Eqn. 6, the relative increase *m_r_* in *M_t_*, is:

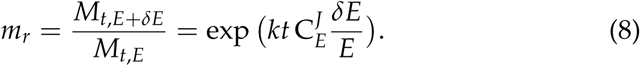

If the doubling time of the system is *t_d_*, then *kt_d_* = *ln*(2) = 0.693 and so the expression for the yield after *n* doubling times is:

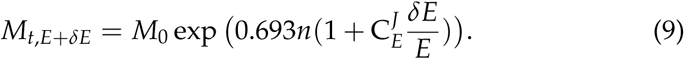

## 4 Alternative derivation

The expression for the perturbed value of *M*_*t*,*E*+*δE*_ can also be obtained more directly without passing via the response coefficient. From Eqn. 3, we can write that the change in *E* produces a change in the rate constant *k* given by:

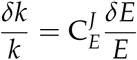

or

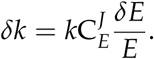

The new value of *k* + *δk* can be substituted directly in Eqn. 4 to give Eqn. 7.

## 5 Response to substantial over–expression of an enzyme

The expressions derived above are, as is generally the case for MCA, only strictly valid for small perturbations about the steady state. Typically, experiments on the effects of over–expression of an enzyme involve substantial changes brought about by insertion of additional copies of the gene for the enzyme and/or changing its promoter to drive higher levels of expression. As noted in the Introduction, although there is no general analytical theory that allows exact calculation of the consequences of such experiments on metabolic fluxes, the consequences of substantial degrees of over–expression can calculated with the finite difference formula derived by Small and Kacser (1993a,b). This states that if an enzyme is over– expressed to *r* times its original activity, the relative increase in flux *f* will be given by:

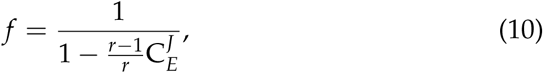

where 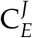 is the control coefficient at the original enzyme level.

In the terminology of this paper, for a finite change in *E* of Δ*E*, *r* = (*E* + Δ*E*)/*E*, and the resulting change in rate constant is (*k* + Δ*k*)/*k* = *f*. Substituting *f k* for *k* in Eqn. 4 gives:

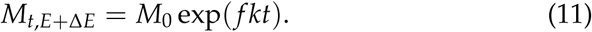

The equation corresponding to Eqn. 8 becomes:

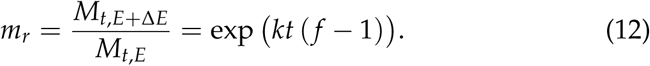

The value of *m_r_* represents the relative increase in yield of product given the flux change *f* induced by over–expression of the enzyme. Note that although *f* is a constant factor of increase in the flux between the control and modified fluxes, the change in yield increases with time, and the combination of Eqns. 10 and 12 allow prediction of the effect on yield of over– expressing an enzyme of known flux control coefficient.

Conversely, given measured values for *M*_0_ and *M*_*t*,*E*_, the value of *kt* can be calculated from Eqn. 1, and then, given *M*_*t*,*E*+Δ*E*_, *f* can be obtained from either of the two preceding equations. Finally, if the ratio *r* of the enzyme activities in the control and over-expressed lines has been measured, rearrangement of Eqn. 10 gives the value of the enzyme’s control coefficient in the control:

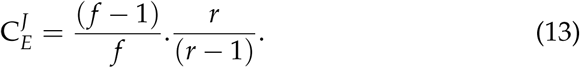

Since Eqn. 10 applies equally to enzyme attenuation, the over-expressed state can be regarded as the reference, with *r*′ = 1/*r* and *f*′ = 1/*f*. Substituting these values in Eqn. 13 gives:

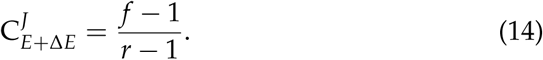

These two equations permit the estimation of an enzyme’s flux control coefficient from experiments where the system is expanding exponentially and the product yields have been measured at two points along the time course without direct determination of the metabolic flux.

## 6 Examples

The potential significant impact of exponential growth on the yield from an engineered organism can be illustrated by plotting the above functions. Fig. 1 illustrates the continuous growth of the relative yield with the length of the exponential phase. By two doubling times, the yield change is already greater than the flux change induced by enzyme over-expression.

**Figure 1:**
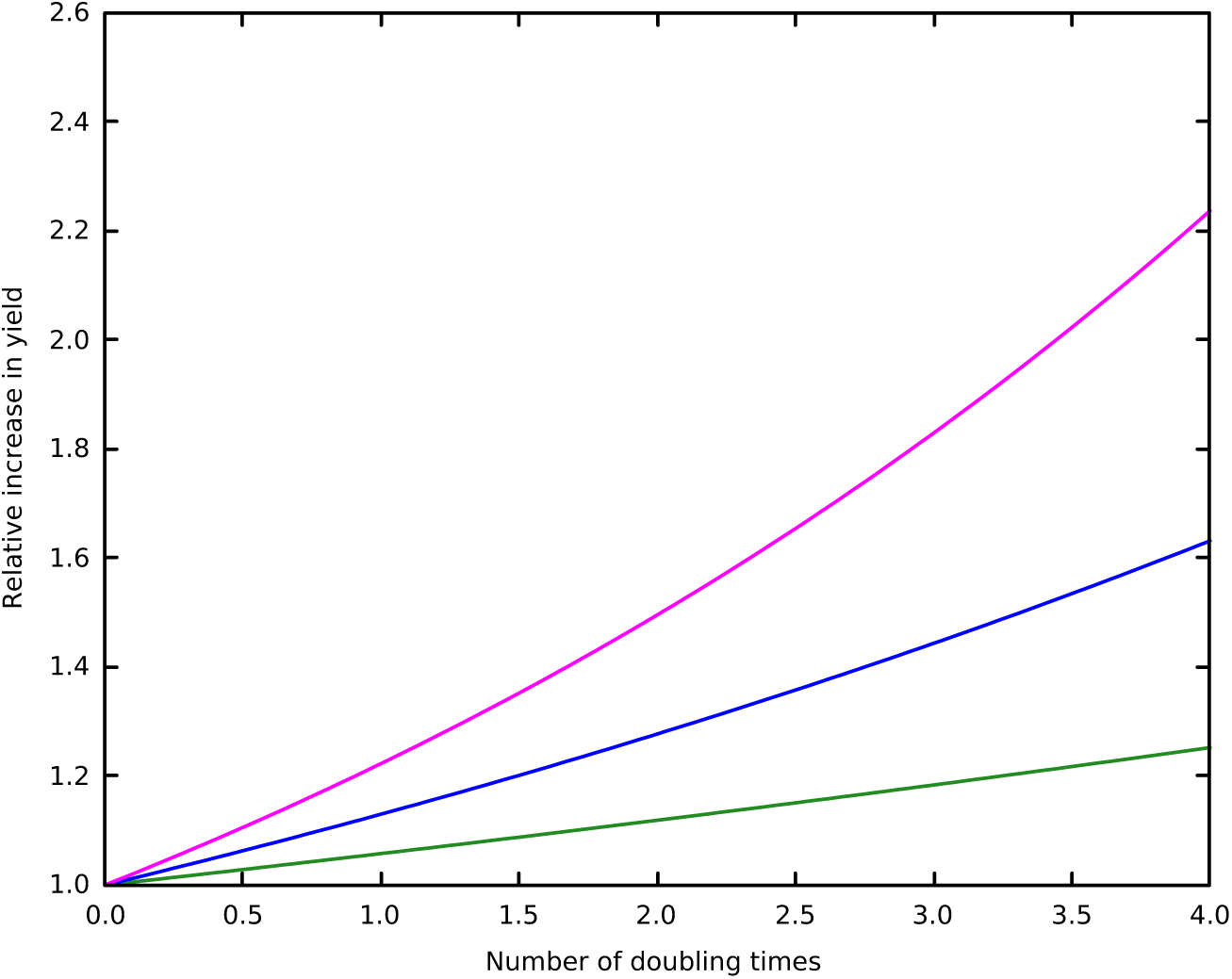
Relative increase in yield by over-expression. Curves are calculated for a two-fold over–expression of enzymes with flux control coefficients of 0.15 (green), 0.3 (blue) and 0.45 (magenta). The corresponding relative increases in flux are: 1.08, 1.18 and 1.29 respectively.

However, as with the effect of enzyme over–expression on flux, the effect on yield becomes less marked as the degree of over–expression increases, as illustrated in Fig. 2, suggesting that it will not usually be worthwhile to pursue very high degrees of enzyme amplification.

**Figure 2:**
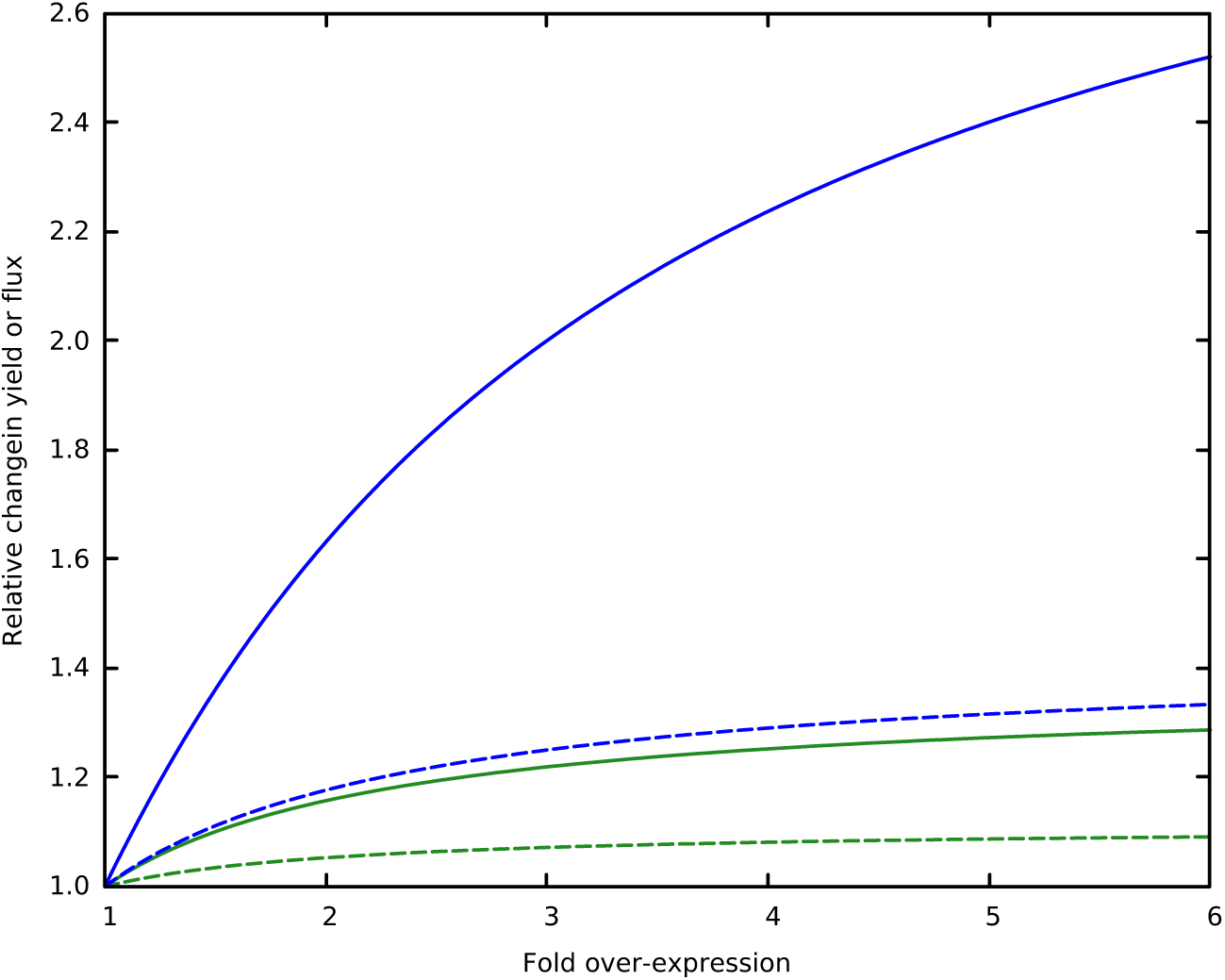
Changes in yield and flux for differenet degrees of over– expression. Relative changes in flux (dashed lines) and yield at 4 doubling times (solid lines) are shown for variable degrees of over-expression of enzymes with flux control coefficients of 0.1 (green) and 0.3 (blue).

## 7 Conclusions

This work has aimed to utilise insights from MCA to applications in metabolic engineering. It builds on the earlier work by Small and Kacser (1993a,b) that related the effects of large changes in enzyme expression on the resulting metabolic flux and extends it to the effects on product yield. The simplest instance of this will be where the desired product is itself biomass or a biomass constituent. However, it will also be relevant to products that are formed in proportion to the biomass (so that the flux to product per unit biomass remains constant throughout the exponential growth phase).

A potential limitation is that it may be unrealistic to expect to manipulate a higher flux to product without a negative impact on the underlying growth rate. This may be possible if the increase in desired product is obtained at the expense of an unwanted by–product rather than of biomass. However, if the increase in flux to product does impact on growth rate or length of the exponential phase, it would be trivial to modify the expressions above to use a different *kt* term for the control and engineered cases, and determine the resulting trade–off between flux to product and growth on the overall yield.

## 8 Acknowledgement

This analysis was motivated by discussions with John Harwood (Cardiff University) and Tony Fawcett (Durham University) about triglyceride accumulation during seed development in oil-seed rape.

## References

Acerenza, L., Sauro, H. M. and Kacser, H. (1989). Control Analysis of Time–Dependent Metabolic Systems, J. Theor. Biol. 137: 423–444.

Fell, D. A. (1997). Understanding the Control of Metabolism, Portland Press, London.

Heinrich, R. and Rapoport, T. A. (1974). A Linear Steady–State Treatment of Enzymatic Chains; General Properties, Control and Effector Strength, Eur. J. Biochem. 42: 89–95.

Kacser, H. and Burns, J. A. (1973). The Control of Flux, Symp. Soc. Exp. Biol. 27: 65–104. Reprinted in Biochem. Soc. Trans. 23, 341–366, 1995.

Meléndez-Hevia, E., Torres, N. V., Sicilia, J. and Kacser, H. (1990). Control Analysis of Transition Times in Metabolic Systems., Biochem. J. 265: 195–202.

Small, J. R. and Kacser, H. (1993a). Responses of Metabolic Systems to Large Changes in Enzyme Activities and Effectors. 1. The Linear Treatment of Unbranched Chains., Eur. J. Biochem. 213: 613–624.

Small, J. R. and Kacser, H. (1993b). Responses of Metabolic Sytems to Large Changes in Enzyme Activities and Effectors. 2. The Linear Treatment of Branched Pathways and Metabolite Concentrations., Eur. J. Biochem. 213: 625–640.

